# Nanoscale mechanical stimulation method for quantifying *C. elegans* mechanosensory behavior and memory

**DOI:** 10.1101/047431

**Authors:** Takuma Sugi, Etsuko Okumura, Kaori Kiso, Ryuji Igarashi

## Abstract

Here, we establish a novel economic system to quantify *C. elegans* mechanosensory behavior and memory by a controllable nanoscale mechanical stimulation. Using piezoelectric sheet speaker, we can flexibly change the vibration properties at a nanoscale displacement level and quantify behavioral responses and memory under the control of each vibration property. This system will facilitate understanding of physiological aspects of *C. elegans* mechanosensory behavior and memory.

## Introduction

Mechanical forces such as touch, vibration and gravity make vital influences on development and homeostasis of living organisms^1,2^. Animals cope with these stimuli by modifying behavior and thereby achieve physical interactions with environment^3^. The mechanism underlying mechanosensory behavior has been studied at various model organisms from bacteria and archaea to mammals^4,5^. However, physiological aspects of those behaviors, such as force and duration enough to sense, are not well understood.

Mechanosensory behavior in *C. elegans* is a traditional paradigm in which to examine mechanism underlying a response to mechanical forces^6–8^. Worms usually respond to nonlocalized vibrations such as the tapping of a cultivated Petri plate with a reversal escape response^7^. In addition, after spaced training for repeated mechanical tap stimulation, worms can habituate to the stimulus and exhibit a decrease in the magnitude of the withdrawal escape response. Therefore, worms can alter behavior based on their past experience^9^. Because the neural circuit underlying this behavior was completely determined, we can easily investigate neural and molecular mechanisms using *C. elegans* cell-specific genetic methods^9,10^. In general, the choice of technique to impose mechanical stimulation and to readout behavioral output is critical for understanding physiology of animal behavior. In *C. elegans*, mechanical stimulation has been provided by tapping only a single plate using a solenoid tapper^11^ or a ROBO cylinder^10^ or by the manual box-drop method^9^, in which a plastic box containing multiple Petri plates is manually dropped onto a hard surface from a constant height. However, in these methods, it has been difficult to precisely change the vibration properties such as frequency, amplitude and duration.

At a cellular level experiment, several mechanical stimulation methods have been established for the study of mechanotransduction^12^. These methods include mechano-clamp using piezo-driven system^13,14^, surface elongation of a flexible silicone elastomer^15,16^, and force application by magnetic particles^17–19^. In addition to these spatially confined methods, Nikukar et al. have recently reported an interesting method, “nanokicking”, using piezo actuator and demonstrated to evoke nanoscale nonlocalized vibration with high frequency (up to 1 kHz) across the entire surface of the Petri plates^20^^,^^21^. However, in this strategy, it was not demonstrated to evoke the frequency above 1 kHz. Furthermore, this method has not been applied to a living animal level experiment, particularly to it’s behavioral experiments.

In this study, we designed an economic nonlocalized vibration device using piezoelectric sheet speaker. This device allows for elaborately changing the vibration parameters and setting these parameters in a desired temporal pattern. Using this device, we clearly quantified reversal responses and memory of worms for nonlocalized vibration. Thus, our new device will facilitate understanding of physiological aspects *C. elegans* mechanosensory behavior by titrating vibration properties in the future.

We previously established the tap stimulation system using the cylinder and actuator, because this method was successfully applied to the quantification of tap habituation behavior in other groups^10,11^. In this system, amplitude of the nonlocalized vibration could be roughly changed, whereas its frequency and duration could not. Therefore, instead of the previous system, we used a piezoelectric speaker to evoke nonlocalized vibration on an agar surface of an NGM plate (Fig. 1 and 2A). The NGM plate on which worms were cultivated (Fig. 1A Step 1) was placed on a circular-shaped actuator of a piezoelectric speaker (Fig. 1A Step 2, and Fig. 2B-D). The actuator was connected to an amplifier (Fig. 2E). This device was also connected to an earphone jack of a desktop computer through an earphone splitter. The earphone splitter enables us to evoke nonlocalized vibration to multiple NGM plates at the same time.

**Fig. 1.**
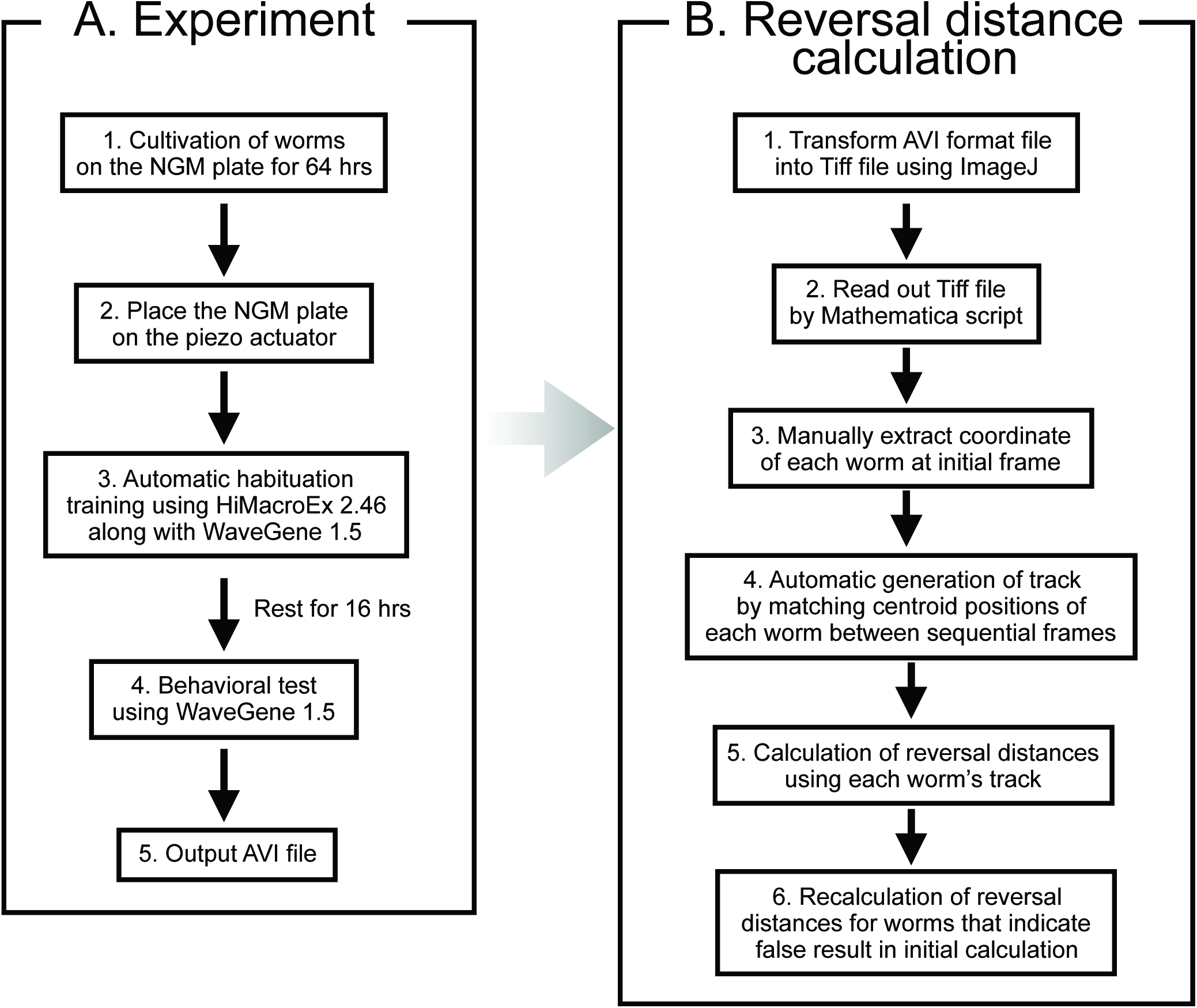
Flowchart for quantification of mechanosensory behavior and memory using piezoelectric speaker in *C. elegans*. (A) Experimental procedure. The flowchart is indicated for examining mechanosensory memory. In the case of only examining behavioral response to mechanical stimulus, step 3 could be skipped. (B) Calculation procedure. The detail instruction is described in Supplementary file.

**Fig. 2.**
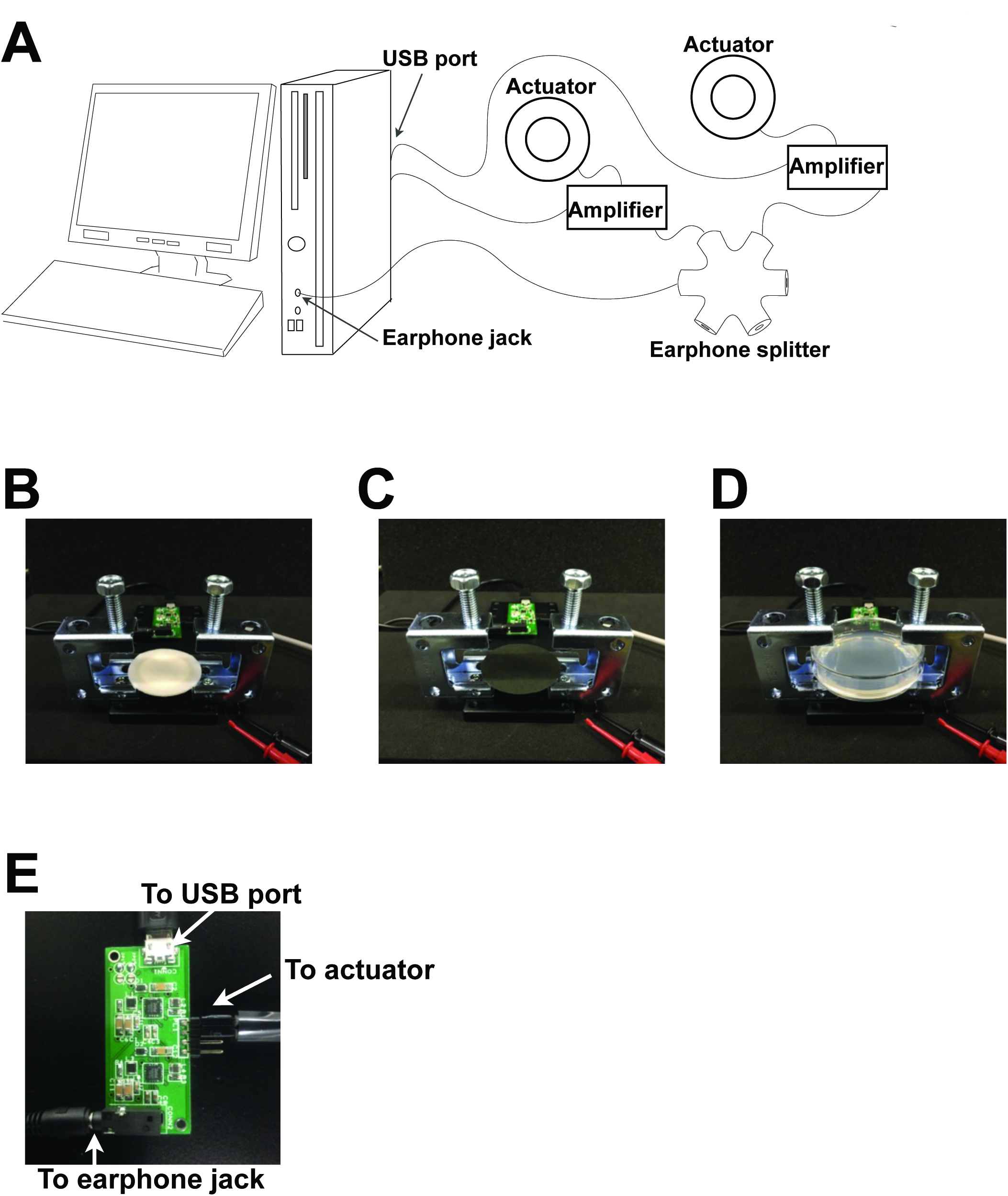
Design of a new nonlocalized stimulation device. (A) Schematic overview of the new device. (B-D) Photos of an actuator without (B) and with a black sheet (C), and with both the black sheet and a Petri plate (D). A black sheet is required for enhancing the contrast between worms and background in each acquired image. The petri plate was fixed by the two screws. Habituation training was conducted inside an incubator and simultaneously applied for the five plates. (E) Amplifier connected to an actuator and a computer.

Amplitude of vibration can be set by volume control of a computer, and this sound volume level was changed in the range of 0 to 100%. On the other hand, the free download software was used for setting the frequency and duration of the vibration. The frequency of vibration can be changed in the range of 0 to 5 kHz. The minimum duration of stimulation is 0.5 sec. Moreover, this devise potentially enables us to generate various waveforms such as sine wave, square wave, pulse wave and white noise.

Furthermore, semi-automation of training of worms was also needed for quantifying habituation memory due to its laborious protocol. The conventional training protocol consisted of five blocks of 20 mechanical stimuli (60 sec interstimulus interval) with a 1 hr rest period between each block^22^. To automatically train large populations of worms, we used a mouse macro system that enables us to program the automatic mouse cursor movement on the computer screen (Fig. 1A Step 3 and Supplementary Fig. 1). To accurately examine habituation memory of trained worms, we have simultaneously prepared untrained worms as a control.

We used laser interferometry to quantify mechanical stimuli evoked by our piezoelectric speaker system. We played 440 Hz sound (the standard tuning frequency) on a piezoelectric speaker and validated the vibration on the center of an agar surface in a Petri plate (Fig. 3A). Expectedly, as shown in Fig. Fig 3B, we succeeded in detecting 440 Hz frequency. Therefore, we have changed the vibration frequency on the center of agar surfaces using WaveGene (Fig. 3C). All the vibration frequencies determined by WaveGene (minimum frequency, 250 Hz) were clearly detected by the vibrometer. In addition, 80 Hz vibration was correctly detected by the accelerometer. These results indicate that our piezoelectric speaker system allows for evoking vibration with accurate frequency on the center of agar surface.

**Fig. 3.**
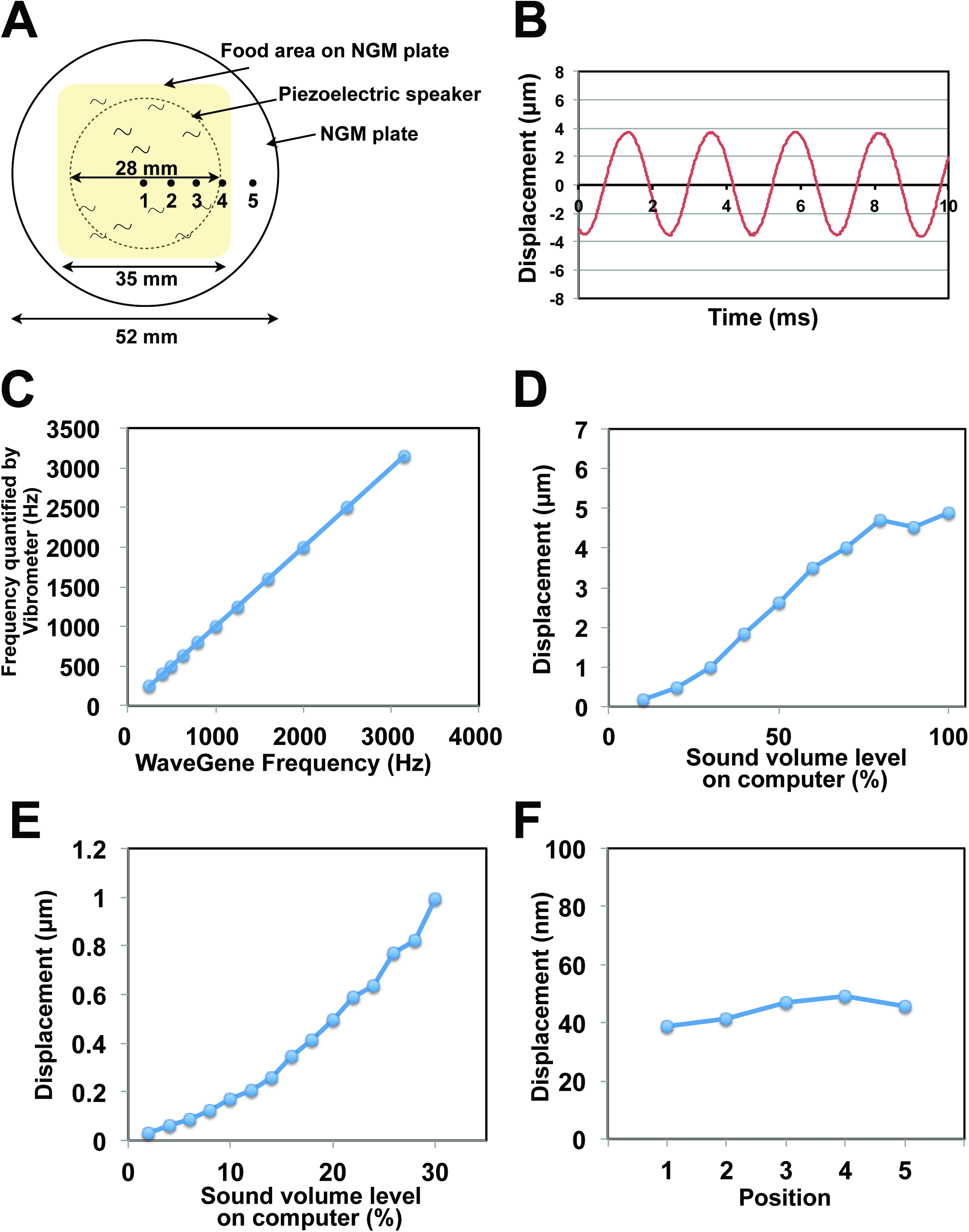
Validation of the new device by laser interferometry. (A) Schematic of the Petri plate, the piezoelectric speaker, and food area on the agar surface. The speaker was represented by dashed line. Displacement was quantified at the indicated positions (1 to 5) as described in Fig. 3 D (B) Detection of the standard tuning frequency (440 Hz) by laser interferometry. The laser was focused on the center of an agar surface. (C) Quantification of various frequencies of vibrations evoked by the new device. The frequency of the evoked vibration was changed using WaveGene, and quantifications were performed by laser interferometry. The computer sound level was set as 50%. (D) Quantifications of the displacements induced by 1kHz vibration in the sound level range of 0 to 100%. (E) Detailed quantifications of the displacements induced by 1kHz vibration, in which the computer sound level was changed in the range of 0 to 20% by 2%. (F) Quantifications of the displacements at the several positions on the agar surface. 1 kHz vibration (computer sound level, 50%) was evoked by WaveGene. Numbers represent distances from the center (1 = center, 2 = 5 mm from center, 3 = 10 mm from center, 4 = 15 mm from center, 5 = 20 mm from center).

We have changed amplitude of vibration by volume control of computer. As shown in Fig. 3D, the amplitude quantified by vibrometer on an agar surface linearly increased as the sound volume level increased, and reached a plateau at 80% of the computer sound level. Importantly, we could detect 1 kHz vibrations with 10% and 20% of the computer sound level as the nanometer scale displacements. Therefore, we next changed the sound level in the range of 0 to 30% by 2% and quantified the displacement in each sound level (Fig. 3E). We could detect the 30.4 nm displacement in 2% of the sound level, and the displacement increased within a nanometer range as the sound volume level became higher. These results reveal that our economic piezoelectric speaker system allows for changing amplitude of vibration at a nanoscale resolution.

Then, we tried to measure 1 kHz vibration (50% of the computer sound level) evoked by WaveGene at the several positions on the agar surface (Fig. 3A and 3F). As a result, we detected almost no frequency deviations across the surface of the Petri plate. These results indicate that our piezoelectric speaker enables us to stimulate almost all worms under the same vibration properties, regardless of their positions on an agar surface.

We prepared both untrained and trained worms and quantified their mean reversal distances at 16 hrs after habituation training (Fig. 1A Step 4-5, Fig. 1B, Fig. 3). Untrained worms exhibited reversal responses to vibration with 1 kHz of frequency, 4.9 μm of amplitude (computer sound level, 100%) and 1 sec of duration (Fig. 3A and 3C). The mean (±SEM) reversal distance of these untrained worms (N = 110) in response to vibration was 1.82 mm (±0.10 mm) (Fig. 3C). On the other hand, in our previous tapping method, mean reversal distance of worms (N = 98) was 1.07 mm (±0.06 mm) (Fig. 3C), which was comparable with that reported in the other group’s paper^11^. Therefore, these quantifications have indicated that worms stimulated by our new system could exhibit longer reversal distances than those stimulated by the old tapping system.

We further examined habituation memory of trained worms (Fig. 3B). The mean reversal distance of the trained worms (N = 135) in response to vibration was 1.36 mm (±0.09 mm). This result revealed that worms trained by our new system showed significantly reduced reversal distance compared with untrained worms (Fig. 3C). Thus, our new system could clearly induce mechanosensory memory through habituation training.

In summary, our system allows for not only quantification of mechanosensory behavior but also training of worms and quantifications of their memory. One of the important advantages for the researcher is that the new system with a device to quantify stimulus is an economic setup (< approximately 130 dollars / a single vibration device) and easily replicated in other laboratories. In addition, we can easily change the vibration properties. Our behavioral experiments have proved the capability of this system. In addition, this system is so compact as to be integrated into another experimental device such as a calcium imaging system. Therefore, our new system will facilitate investigation of physiological aspects of behavior and neural circuitry in the future.

## Online Methods

### Strain preparation

We used wild-type N2 Bristol strain for all behavioral experiments. This strain has been maintained and handled using standard methods^23^.

### Vibration device construction and it’s validation

To control properties of nonlocalized vibration precisely, a new system was constructed using the piezoelectric sheet speaker (THRIVE, pzBAZZ μSpeaker B35) as an actuator and the amplifier module (THRIVE, 0530AMPZ) connected to a computer earphone jack via an earphone splitter. The diameter of sheet speaker was 42 mm, and frequency could be increased at least up to 5 kHz. The amplitude could be also changed by volume control of the computer in the range of 0 to 100% (Dell PRECISION T1650 desktop computer). The minimum duration of vibration is 0.5 sec. The mechanical stimuli were quantified by laser vibrometer V100 (Denshigiken Corp.) with PicoScope oscilloscope.

### Behavioral recording

All worms’ behaviors were recorded at > 7.0 frames / sec using USB-controlled CCD cameras (Sentech, STCTB83USB-AS), which were each coupled to a 25 mm focal-length C-mount machine vision lens (Azure, AZURE-2514MM) and C-mount adaptor (5 mm thickness, 30CMA-R). Each pixel in the captured images corresponds to a 25.4 μm × 25.4 μm area in each Petri plate. The total field captured was 26.0 mm × 19.5 mm.

### Software

Two free download software were used for automatic stimulation; WaveGene Ver. 1.50 for control of vibration properties and mouse macro HiMacroEX 2.46 for automatic habituation instead of manual operation. The software was compiled and bench marked on a Dell PRECISION T1650 desktop computer (Dell). The script written in HiMacroEx 2.46 was indicated in the Supplementary Fig. 1.

The reversal distance was calculated according to a previously reported method^10^. At first, an acquired AVI format movie was transformed into Tiff-format sequential images using ImageJ software (NIH). This sequential image file was used for subsequent motion analysis of each worm. Image-processing software for the quantification of each worm’s reversal distance was written in Mathematica 9.0 (Wolfram). The coordinate of each worm at initial frame was extracted manually. Then, the centroid of each worm was calculated and a track was generated by matching centroid positions between sequential frames. This track was used to calculate reversal distance. Initially, reversal distance of each worm was calculated automatically by the software. After this initial calculation, a frame number in which reversal movement was completed was manually confirmed for each worm using output result. Then, correct frame numbers for worms that indicate incorrect frame numbers in the initial calculation was extracted manually and put into the software for recalculation of their reversal distances. The detail instruction for this software was described in the Supplemental file.

### Habituation training

Worms were cultivated on 60 mm Petri plates (Thermo Fisher Scientific, #150288) containing 10 mL of NGM with 2% agar, on which *Escherichia coli* OP50 was seeded. On the first day, 8-9 worms were deposited onto each NGM plate and cultivated at 20 °C. After 3 hrs, the deposited P0 worms were removed to segregate the F1 progeny. The progeny were cultivated for 64 hrs at 20 °C. An NGM plate on which worms were cultivated was placed on an actuator of a piezoelectric sheet speaker and stimulated through WaveGene. The training protocol was flexibly customized using Mouse macro HiMacroEX 2.46. In this study, the conventional training^22^ was adopted for worms within a 20 °C incubator (ADVANTEC). The nonlocalized vibrations were evoked at every 1 min for 20 times, and this stimulation sequence was repeated five times with a 1 hr interval.

**Fig. 4.**
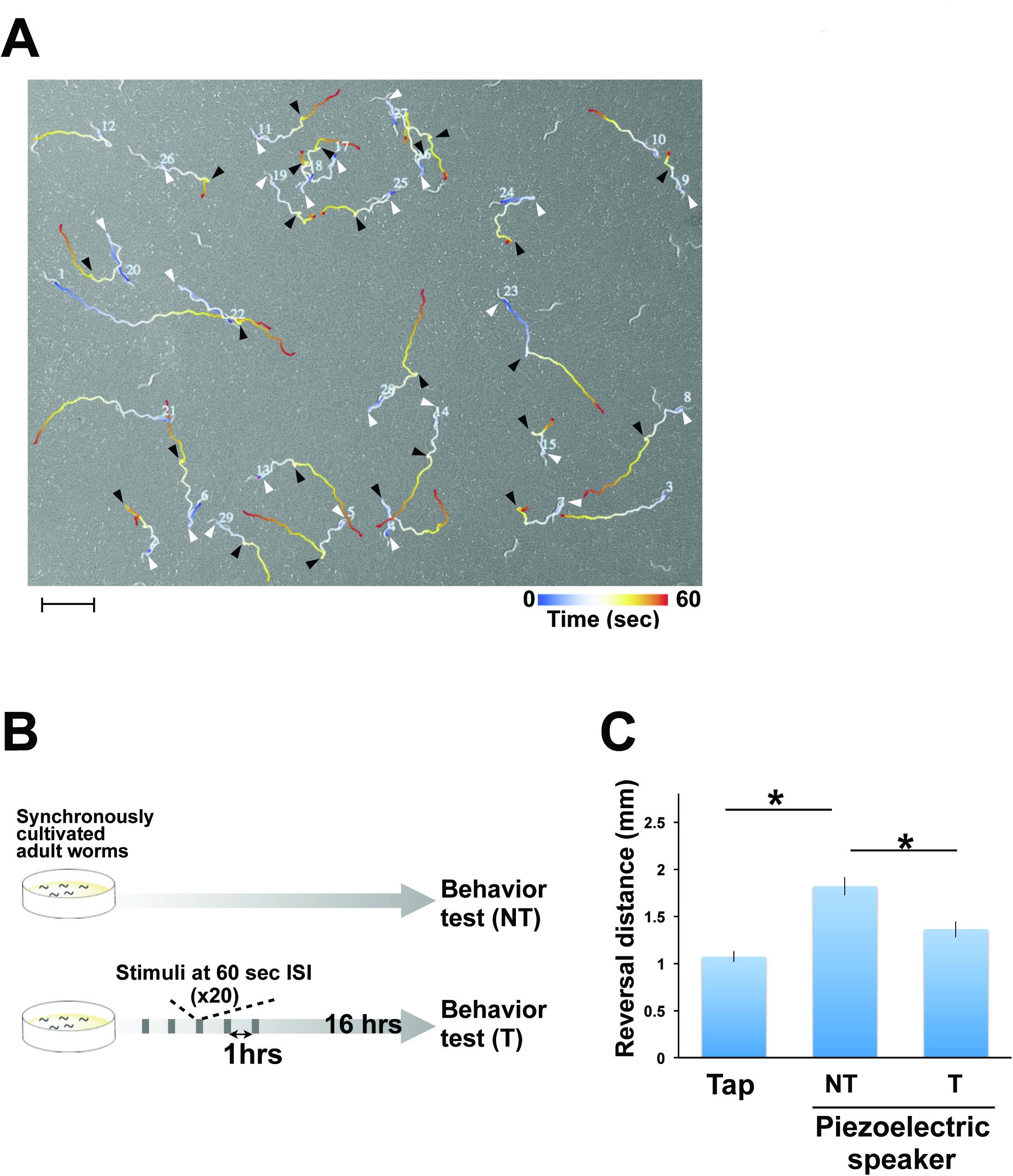
Behavioral experiments. (A) Trajectories of worms’ reversal responses to nonlocalized vibration induced by the new system. The vibration with 630 Hz of frequency, 4.5μm of amplitude (computer sound level, 50%), 1 sec of duration was delivered to the Petri plate at 10 sec after starting behavioral recording. The white and black arrow heads indicate start and end positions of each worm’s reversal movement, respectively. Worms that suddenly accelerated forward movement in response to vibration were not marked by the arrow heads. Scale bar, 2 mm. (B) Scheme of behavioral experiments. The conventional protocol (five blocks of 20 tap stimuli (60 sec interstimulus interval) with a 1 hr rest period between each block) was used for habituation training. Behavioral quantifications were performed at 16 hrs after habituation training. (C) Quantifications of mechanosensory behavior and memory using the old tap system and the new piezoelectric sheet speaker system. Reversal distances of worms that were not trained (NT) and trained with the conventional protocol (T) were quantified at 16 hrs after training. The vibration with 1 kHz of frequency, 4.9 μm of amplitude (computer sound level, 100%), 1 sec of duration was delivered to the Petri plate at 10 sec after starting behavioral recording. More than 80 worms were examined in each experimental condition. Error bars indicate SEMs. Statistical comparisons were performed using t tests. \**P* < 0.01

## Acknowledgements

We thank *Caenorhabditis* Genetic Center for sharing strains. T.S. and R.I. were supported by the Japan Society for the promotion of Science, Japan Science and Technology Agency under Precursory Research for Embryonic Science and Technology (PRESTO). T.S. was supported by Mochida Memorial Foundation for Medical and Pharmaceutical Research.

